# An eco-evolutionary theory of host-associated microbiomes

**DOI:** 10.64898/2026.02.10.705025

**Authors:** Gui Araujo, Torsten Thomas, Nicole S Webster, José M Montoya, Miguel Lurgi

## Abstract

Host-associated microbiomes often display host specificity and heritability, yet the evolutionary processes under which such structured communities first emerge are still unclear. In particular, the conditions by which intergenerational (i.e. vertical) transmission of microbes can evolve and generate host-specific microbiomes are still unresolved. Here, we present an eco-evolutionary theory of microbiome assembly under minimal assumptions of microbial dynamics (i.e. neutrally driven by environmental fluctuations) and host control. We consider the adaptive evolution of microbial and host traits, including microbiome size and vertical transmission. We show that environmental fluctuations can generate enough among-host microbial variation to enable host-level selection favouring beneficial microbiome configurations. Vertical transmission can then evolve and, even when weak, allow microbiome specificity to be inherited and amplified across generations despite continuous influx from the external environment. Selection is most effective at intermediate levels of environmental fluctuation and host lifespan, revealing fundamental trade-offs between stochastic assembly, inheritance, and dispersal of microbes. The resulting microbiomes are dense, host-specific, and heritable, yet retain high intraspecific variability and lack strict phylosymbiosis. Simulated patterns of microbial dominance, diversity, and host-microbiome dissimilarity closely match those observed in nature, as evidenced using marine sponge microbiomes. Our results provide a mechanistic theory for the early evolution of host-associated microbiomes, showing that beneficial and species-specific communities can arise through selection and inheritance prior to the evolution of dedicated host-control mechanisms.

## Introduction

Symbiotic associations with microbial communities are prevalent across animal hosts [1]. These communities often exhibit strong host species specificity [2,3], reflecting processes that tend to benefit the host by improving survival and reproductive success [1]. Microbes may provide these benefits through multiple mechanisms, including aiding digestion [4], producing bioactive compounds [5], protecting against pathogens [6], and modulating the host’s internal environment [7]. Since host survival often enhances microbial persistence and transmission, these interactions can create mutual fitness benefits, even in the absence of reproductive advantages to microbes within the microbiome [8]. Ultimately, these communities emerge from a combination of ecological assembly processes and responses to environmental changes, which can accumulate over evolutionary timescales.

Host-associated microbiomes exhibit similarities to other ecological communities, assembling through processes of colonisation, persistence, and growth [9]. Microbial types are acquired, transmitted, and proliferate through various routes [9], even via concurrent transmission pathways, such as taxa-specific microbial transmission associated with social and environmental routes observed in mice [10]. Microbial transmission is therefore among the foundational processes structuring host-associated microbiomes. Vertical transmission, the transfer of microbes from parent to offspring, promotes community fidelity and heritability by preserving similarity in parent and offspring microbiome compositions [11-14]. In contrast, horizontal transmission, the acquisition of microbes from the environment or other hosts, can erode parent-offspring fidelity and promote microbiome differences across related individuals [15]. Hence, the relative strength of vertical and horizontal transmission is a key determinant of microbiome composition and specificity [11,13,16,17]. Despite this understanding, the evolutionary conditions under which vertical transmission itself emerges and becomes selectively favoured remain poorly resolved.

Theoretical work poses that microbiome assembly processes can be deterministic, thus making microbiome structure predictable; but they can also be strongly influenced by stochastic processes such as those arising from, for example, environmental fluctuations [9,18-20]. Empirical microbial communities have been shown to be sensitive to stochasticity generated by natural variation in abiotic conditions, host physiology, and random environmental perturbations [21,22]. Through synergistic interactions with transmission processes, environmental stochasticity can amplify variation among hosts relative to variation within a host’s internal environment, which is often under stronger deterministic control [9,18]. As a result, the balance between deterministic and stochastic drivers impacts the extent of non-random organisation in microbiome composition, thereby affecting microbiome-mediated host benefits [18]. Fluctuations in population abundances can also act as a source of diversity [23], enabling random shifts in composition while counteracting the convergence imposed by deterministic processes. For example, environmental variability can offset deterministic effects of colonisation order (priority effects) and allow for the establishment of late-arriving species that would otherwise become extinct [24]. Random perturbations in microbial abundances can therefore modulate the arrival and persistence of functionally beneficial microbes and their transmission.

Microbial transmission influencing assembly and inheritance, alongside environmental fluctuations that promote microbial variability, thus emerge as key mechanisms shaping the evolution of microbiomes and the emergence of microbiome-related traits and interactions. Previous theoretical work has shown that vertical transmission and stochasticity can enrich specific microbes, either by boosting their host-specific success that would otherwise be disadvantaged [8,25], or when combined with host-mediated changes in regional microbial availability linked to microbial abundances within hosts [17,26,27]. However, existing models have focused on simple microbiomes with few taxa [8,27] or have examined the consequences of an already established process of vertical transmission [25,26]. As a result, it remains unclear how vertical transmission itself evolves, which mechanistic conditions favour its selection, and whether it is sufficient to generate and maintain structured, diverse microbiomes as both microbes and hosts evolve [28].

To address this gap, we developed a mechanistic theory of microbiome assembly and host-microbe evolution to ask whether vertical transmission can evolve under neutral microbial dynamics (i.e. without advantages in microbial reproductive success) in fluctuating environments. We then identify the mechanisms responsible for the evolutionary emergence of host-microbiome associations in this setting. Our results show that, in fluctuating environments, the presence of beneficial microbes (i.e. microbes that enhance host reproductive success) can drive the evolution of vertical transmission and the emergence of specific microbiomes, even in the presence of horizontal transmission that erodes community fidelity. Our simulations successfully replicate empirical diversity distributions at both individual and species levels, as well as the relationship between microbiome dissimilarity and host phylogenetic distance observed in marine sponges. Together, these findings provide new theoretical insights into how environmental perturbations, transmission, and selection interact in the ecology and evolution of complex symbioses, contributing to the growing need to understand and preserve host-associated microbial communities in an era of rapid environmental change [29-31].

## Results

To explore the evolutionary assembly of microbiomes from relatively simple origins into host-specific communities, we developed a mathematical model that captures the joint ecological and evolutionary dynamics of hosts and their microbes. We conducted simulations in which a host’s microbiome density (i.e. their carrying capacity *K*) evolved from initially small values, and vertical transmission was selected to increase from an initial state of no vertical transmission. We then tested whether distinct microbiome structures could emerge spontaneously, yielding dense, vertically transmitted communities with compositions that contrast with the microbial communities from the external environment. Instead of assembling microbiomes via active enrichment or host-control mechanisms, we assumed that microbes follow neutral ecological dynamics, with equivalent reproductive dispersal rates. This setting allowed any emerging structure to result from host-level eco-evolutionary processes.

The model considers three processes that we hypothesise shape host-associated microbiomes. *Microbiome assembly* describes how microbial populations colonise, grow, compete, and fluctuate within individual hosts. *Host dynamics* links microbiome composition to host fitness, determining the extent to which individual hosts contribute offspring to the next generation. Finally, *trait evolution* allows both hosts and microbes to change (via mutation) and evolve via natural selection. We define traits as evolvable parameters representing biological attributes with meaningful effects on reproductive fitness. In this case, we consider only mutations that directly lead to phenotypic changes. Hosts evolve through changes in the magnitude of vertical transmission, density of the microbial community they carry (i.e. *K*), and metabolic complementarity with microbes (a trait-matching of hosts to beneficial microbes), and microbes through changes in their own metabolic complementarity trait.

### Microbiome assembly

Microbiome assembly was modelled as a stochastic ecological process in which each host *h* contains *S* microbial types, the *i*-th type with abundance *x*_*ih*_. All *S* microbe types are always existing in the external environment and enter into hosts in a constant total influx. The process of assembly plays out as microbes increase their abundances until approaching the host’s limit (see example in Fig. 1A). The dynamics of microbial growth inside hosts follow a stochastic differential equation:

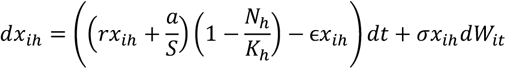

where *N*_*h*_ = ∑_*i*_ *x*_*ih*_ is the total microbiome abundance. This formulation captures how microbial populations grow within a finite host carrying capacity *K*_*h*_, while simultaneously experiencing turnover (*ϵ*) and environmental noise (*σ*). The first term combines two neutral processes: intrinsic growth at rate *r* and microbial immigration from the environment following a total influx *a*, equally divided among all microbial types. The stochastic term represents random environmental fluctuations, modelled with normally distributed increments *dW*_*it*_ ∼ 𝒩(0, *dt*). As the microbiome grows, the space within the host fills (the total abundance *N*_*h*_ tends to *K*_*h*_), and growth slows down asymptotically. Microbiome assembly is illustrated in Fig. 1A.

**Fig. 1.**
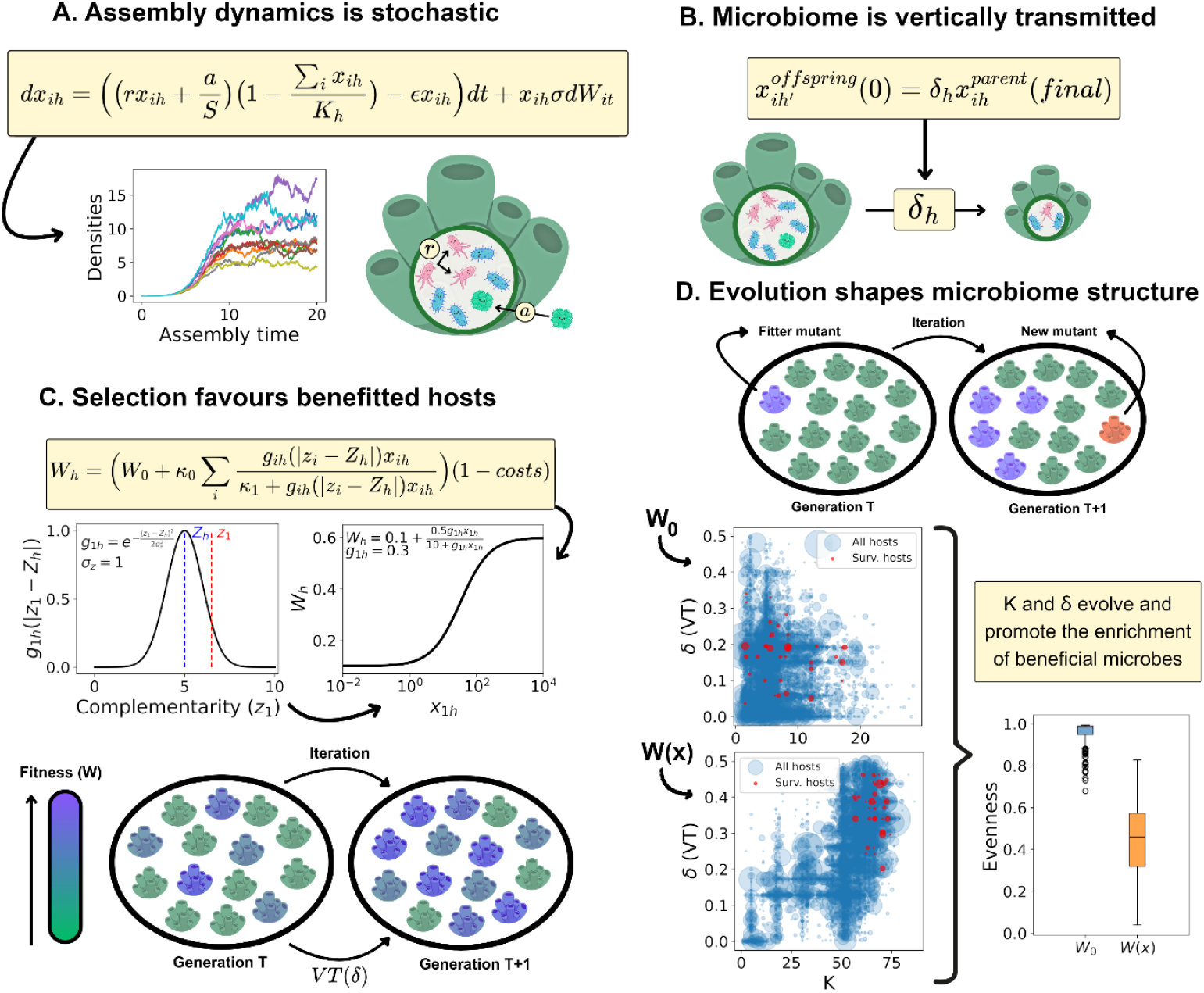
Mechanisms of the eco-evolutionary model and the emergence of specific microbiomes. Pipeline of the modelling approach followed in this paper. (A) Microbiomes assemble neutrally within each host under environmental fluctuations (σ). Microbial abundances (or densities), *x*_*ih*_ for the *i* − *th* type (indicated by different colours) within host *h*, rise through intrinsic growth (*r*) and immigration from the environment (horizontal transmission, *a*) until reaching the host’s carrying capacity (*K*_*h*_), after which communities fluctuate around equilibrium and experience constant turnover (ϵ). (B) During host reproduction, offspring inherit a fraction (δ_h_) of the parental microbiome through vertical transmission, which seeds a new assembly process in the next host generation. (C) Host reproductive fitness increases with the abundance of beneficial microbes, following a saturating response determined by the match between host (*Z*_*h*_) and microbial (*z*_*i*_) complementarity traits. This response is modulated by a normally distributed benefit kernel *g*_*ih*_(|z_i_ − *Z*_*h*_|). Both δ_h_ and K_h_ incur proportional maintenance costs to fitness. The example shown illustrates how fitness rises with the abundance of a beneficial microbe type (*x*_1*h*_; g_1h_ = 0.3). With vertical transmission, average fitness is expected to increase across generations as better-adapted hosts reproduce and propagate their microbiome structure. (D) Host traits for vertical transmission (δ_h_), carrying capacity (*K*_*h*_), and complementarity (*Z*_*h*_) evolve under mutation (μ_H_), while microbial complementarity also changes across generations. Starting from no vertical transmission and minimal microbiome size, simulations run for 2,000 generations show that systems lacking microbial benefits (*W*_0_) fail to evolve specific microbiomes, whereas those with beneficial interactions (*W*(*x*)) evolve strong vertical transmission and increased microbiome size. The resulting decrease in evenness indicates the emergence and persistence of non-homogeneous, host-specific microbiomes dominated by beneficial taxa. Surviving hosts were defined as the 50% better-fitted individuals at the end of the last generation, and the boxplot shows the distribution of evenness for these hosts. See *Methods - Simulations* for parameter details.

### Host dynamics

Host populations consist of *H* individuals, each living for a period *T* and carrying an independent microbiome that assembles over their lifetime. At the end of each generation, individual hosts reproduce with probability proportional to their fitness *W*_*h*_, which depends on how much benefit they obtained from their associated microbiome:

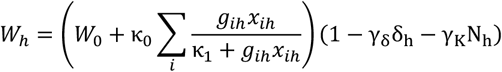

The first term represents benefits from interactions with microbes over a constant baseline *W*_0_. Hosts gain increasing but saturating returns from the total abundance of beneficial microbes, as determined by the benefit due to trait matching function *g*_*ih*_ defined below. The second term introduces costs of transmitting and maintaining the microbiome, weighted by *γ*_*δ*_ and *γ*_*K*_, respectively. Reproduction follows a Wright-Fisher process where offspring are sampled from the parental generation with probabilities proportional to *W*_*h*_. In this process, each iteration features an entirely new generation composed of offspring that replaces individuals from the previous generation, keeping constant the total number of hosts *H*. Each offspring inherits, as their initial microbiome state, a fraction *δ*_*h*_ of its parent’s final microbial community, such that the microbiome composition of the parent is transmitted to the offspring in a smaller total size:

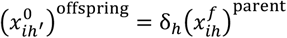

Thus, *δ*_*h*_ represents the degree of vertical transmission, from purely environmental acquisition (*δ*_*h*_ = 0) to partial inheritance (*δ*_*h*_ > 0) of the parental microbiome (Fig. 1B).

### Trait evolution

Both hosts and microbes can evolve via random mutation of a series of traits. Each host species is defined by three heritable traits: carrying capacity *K*_*h*_, vertical transmission *δ*_*h*_, and the metabolic complementarity trait *Z*_*h*_, which may mutate with probability *μ*_*H*_ at reproduction. When a mutation occurs, one trait is randomly chosen to shift its value by a small amount (see Methods - Simulations for details). Microbes possess analogous metabolic complementarity traits *z*_*i*_ (for type *i*), defining how well they match the host’s *Z*_*h*_. The benefit function *g*_*ih*_ depends on the distance between these two traits within a circular trait space of size *ζ*:

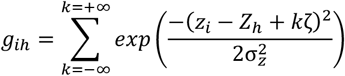

This function ensures that microbial benefit decays smoothly with trait mismatch while respecting periodic boundaries, maintaining *g*_*ih*_ ∈ [0,1] (the whole fitness expression with complementarity is illustrated in Fig. 1C).

Microbial evolution proceeds by creating a new microbial type in the microbial pool with probability *μ*_*m*_ at the end of each host generation, thereby increasing the richness of microbial types *S*. Microbial richness within hosts increases in general with new microbe types, but we considered an abundance threshold of 10^−6^ below which it was set to zero. New microbes inherit their parent’s metabolic complementarity (*z*_*i*_) with a small random deviation. Chances of mutation for each type are proportional to its overall abundance, ensuring that each individual microbe has the same propensity to mutate.

### The evolution of microbiome inheritance and specificity

We initialized the model with *H* = 1,000 identical hosts assembling rudimentary microbiomes from an initially empty state (all 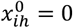), starting with five microbial types (*S* = 5), small carrying capacity (*K*_0_ = 5), no vertical transmission (*δ*_0_ = 0), and random metabolic complementarity trait (i.e. hosts lacked any specific or heritable microbiome). Over 2,000 generations, however, average host fitness rose steadily before stabilizing (Fig. S1). This increase in fitness results from enriching beneficial microbial types. Simulations exhibited a joint evolution of larger host carrying capacities and stronger vertical transmission, ensuring that dense beneficial communities were inherited and made specific across generations (Fig. 1D, replicates in Fig. S2)

To assess the impact of microbial benefits on the evolution of host traits and emergence of community compositions with enriched beneficial types, we ran simulations without any benefits or fitness distinctions (*W*(*x*) = *W*_0_). In this case, hosts failed to expand (i.e. increase *K*) or differentiate their microbiomes, and communities remained compositionally homogeneous, as the external environment, with high evenness (Fig. 1D). This confirms that, in the absence of any host-control mechanism and under environmental fluctuations, selection of hosts receiving more microbial benefits alone was sufficient to drive the evolution of vertical transmission and the emergence of specific, heritable microbiomes.

### Mechanisms driving microbiome evolution

To understand the mechanisms underlying the evolution of specific microbiomes, we analysed how the processes of microbial assembly and host selection interacted to generate heritable communities with specific structures. A key quantity emerging from the assembly model is the relative strength of microbial immigration versus intrinsic growth, expressed by the parameter 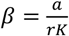. This ratio captures how strongly the microbiome depends on external influx relative to self-replication within the host.

Using the deterministic version of the model (*σ* = 0), we identified a separation of timescales between fast microbial assembly and slow turnover toward the equilibrium state (Fig. S3). From this, we derived a characteristic assembly timescale that sets a lower bound for the host’s entire lifespan *T* that enables a full microbiome assembly (Fig. S4). Within this framework, we obtained an analytical expression for a *structure retention factor, C*, describing how much the abundance differences among any two microbial types are preserved across generations (see Supplementary Material for analytical derivation of this expression):

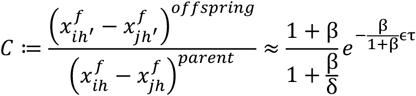

The parameter τ is the additional host lifespan after assembly. We necessarily have that *C* < 1, indicating an inevitable erosion of structure, with *C* = 1 representing an unattainable case of perfect inheritance. Exploring *C* across the parameter space (Fig. 2A) revealed that short lifespans (small τ) and low horizontal transmission relative to intrinsic growth (small *β*) promote the maintenance of microbiome structure. Interestingly, even weak vertical transmission (small δ) was sufficient to maintain substantial structure in microbiomes, as most gains in *C* occur at small *δ*. In contrast, long host lifespans imposed strict upper bounds on maintenance, causing community homogenization over time. The effect of *C* < 1 is evident in scenarios where diverse initial microbiomes converge to a uniform composition over successive generations despite high vertical transmission (Fig. 2B).

**Fig. 2.**
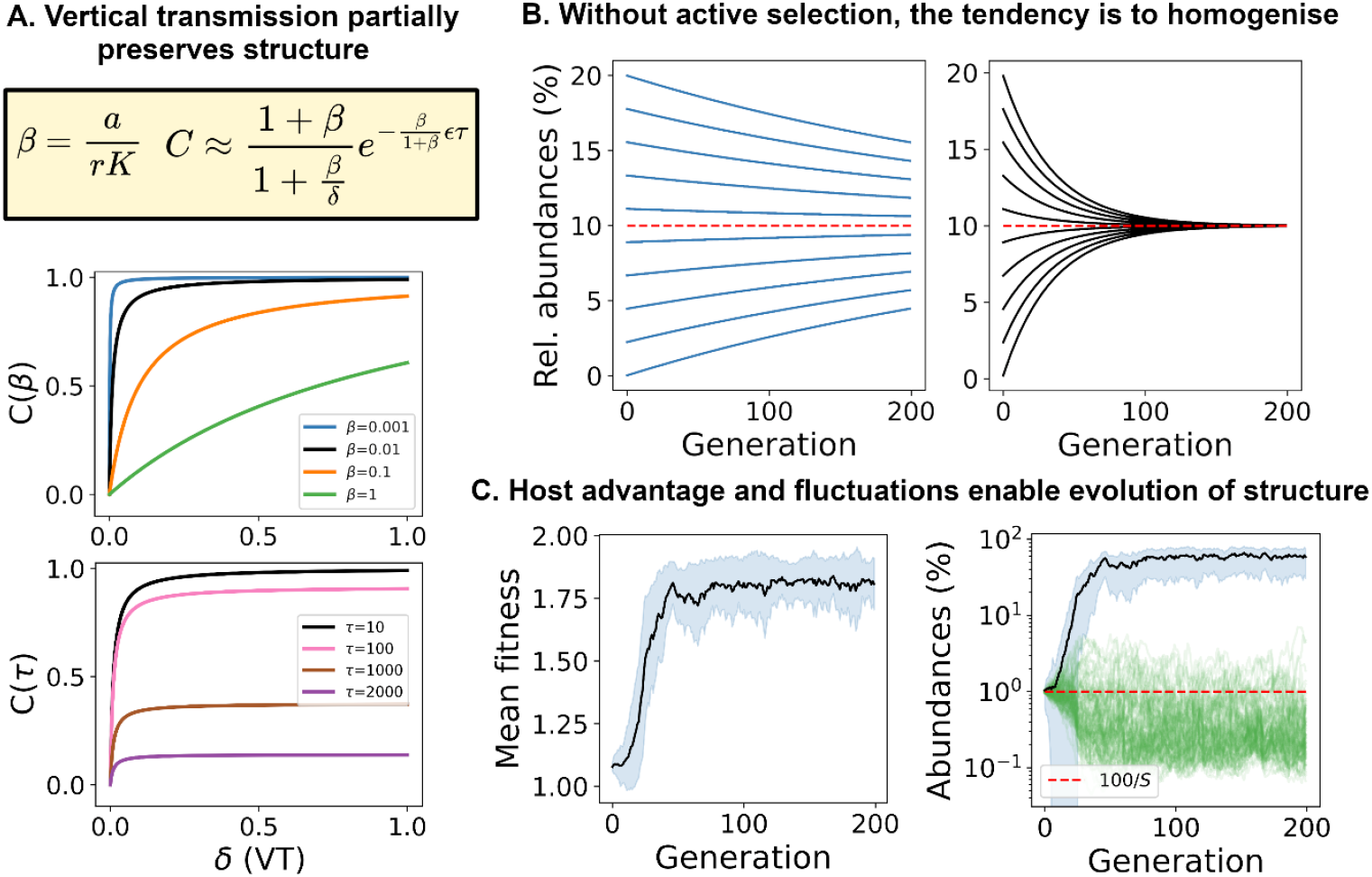
Vertical transmission enables selection in fluctuating environments. Analytical exploration of the deterministic system (without fluctuations) reveals key mechanistic relationships underlying microbiome assembly. (A) The relative influence of microbial immigration and intrinsic growth, expressed as 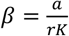, together with host lifespan after assembly (τ) and vertical transmission (δ), determines the factor of microbiome structure maintenance across host generations (*C*). This factor quantifies how strongly differences in microbial abundances are preserved as vertically transmitted microbes are diluted by horizontal transmission. Plots show *C* as a function of δ for different values of β (top, τ = 10) and τ (bottom, β = 0.01), with *r* = 1 and ϵ = 0.1. *C* never reaches 1 (perfect maintenance), and most of its increase occurs at small δ, indicating that even weak vertical transmission substantially improves inheritance of community structure. (B) Vertical transmission partially preserves abundance differences across 200 host generations for different values of *C*(δ), with *δ* = 0.4, τ = 10, ϵ = 0.1, and β = 0.001 on the left, and β = 0.01 on the right illustrating a complete collapse to homogeneous communities of abundances 100/S (red line). (C) When environmental fluctuations and host benefits by associated microbes are considered, the dynamics shift. Selection together with vertical transmission counters the homogenising effects of horizontal transmission. In these simulations (100 hosts, *K* = 100, δ = 0.4), a single beneficial microbe among 100 species is rapidly favoured, increasing both average host fitness (left) and the average relative abundance of the beneficial microbe (right, in black). Averages are calculated for the 100 hosts right after assembly (*T* = 10), standard deviations are shown in shaded blue (very large in the beginning) and homogeneity is represented by the red line. *See Methods - Simulations* for parameter details.

Selection acting through differential host reproduction driven by beneficial microbes can, in principle, counteract this homogenizing tendency. Yet, for selection to operate, there must first be variation to act upon. Environmental fluctuations generate this variation, producing hosts that, by chance, favour the proliferation of beneficial microbes. When vertical transmission is present, these favourable configurations are preferentially inherited, counteracting the homogenizing drift. In simulations with one beneficial microbe among 100 types (*H* = 100, *β* = 0.001, *T* = 10, *δ* = 0.4) and initially empty communities, measured right after assembly, beneficial microbiomes rapidly evolved and remained consistently structured across hosts (Fig. 2C). Removing vertical transmission, however, led to unstructured communities and no fitness improvement (Fig. S5). Thus, vertical transmission enables effective selection in a fluctuating environment, allowing host-beneficial microbiomes to emerge and persist despite the inherent tendency towards compositional uniformity.

Explorations of the model’s parameter space revealed a broad region in which beneficial microbes were effectively selected, reaching relative abundances above 1% among 100 microbial types. This success was not restricted to perfectly advantageous microbes: even when the beneficial type reproduced more slowly than others (*r*_1_ < 1), selection could remain effective (Fig. 3A). Thus, vertical transmission and host-level selection can sustain beneficial associations despite reproductive disadvantage in the beneficial microbes.

**Fig. 3.**
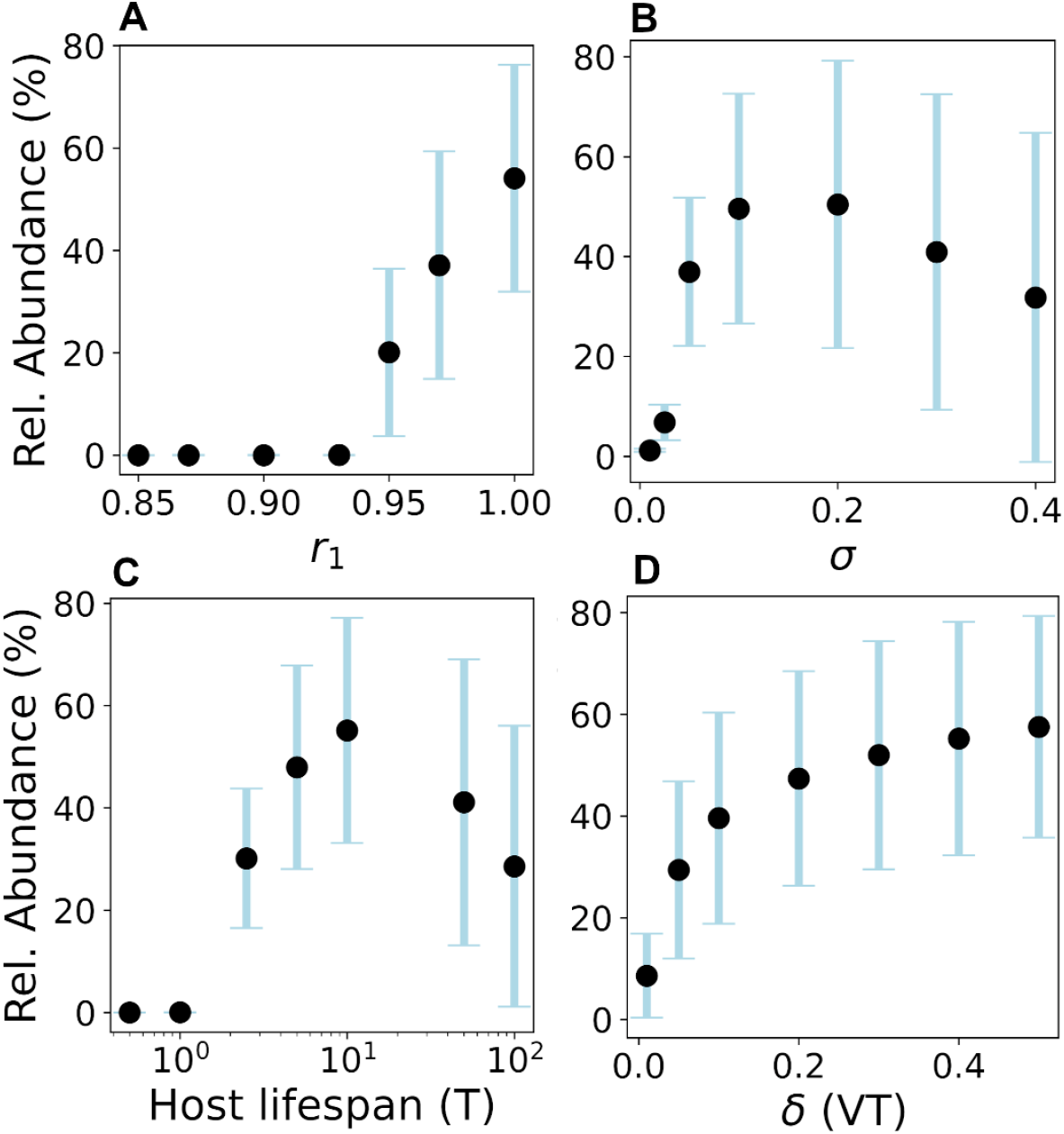
Nonlinear and optimal responses of microbiome evolution to key ecological parameters. (A) Breaking neutrality across the microbes within the microbiome by reducing the reproductive rate of the only beneficial microbe (*x*_1_) while keeping others at *r* = 1 reveals that *x*_1_ can still dominate even when reproductively disadvantaged. (B) Varying the intensity of environmental stochasticity exposes an optimal value. Small amounts of noise fail to generate sufficient variation for selection, whereas excessive noise disrupts structure stability. (C) Host lifespan (T) shows a similar pattern, with selection being strongest at intermediate durations. Short lifespans truncate microbiome assembly, while long lifespans drive homogenisation (as predicted by low C(δ); Fig. 2A). (D) Increasing vertical transmission (δ) enhances selection efficiency, but with diminishing returns, consistent with the shape of the *C*(δ) curves. Simulations follow the host-microbiome dynamics described in Fig. 2C, with one beneficial microbe (*x*_1_) among 100 types and no evolving traits. *H* = 1000 hosts were evolved for 100 generations. Across all plots each point shows the mean (black dots) ± s.d. (blue bars) across all hosts in the last 20 generations. In a homogeneous state, each microbe represents 1% relative abundance; thus, a higher abundance of *x*_1_ indicates successful selection and the emergence of microbiome structure. See *Methods - Simulations* for parameter details.

Environmental stochasticity showed a characteristic *intermediate optimum* (Fig. 3B). When fluctuations are too weak, little diversity emerges for selection to act upon; when too strong, compositional stability in the microbiome is lost. Similarly, host lifespan (*T*) displayed a clear optimum near the characteristic assembly timescale (Fig. 3C): short-lived hosts interrupted the assembly process, while overly long lifespans after assembly allowed stochasticity, turnover, and horizontal influx to erode inherited structure. The strength of vertical transmission (*δ*) followed the expected pattern: low values are already sufficient to promote effective selection, with diminishing returns at higher levels (Fig. 3D). Other parameters behaved consistently with expectations (Fig. S6), reinforcing the argument that microbiome selection stands on a balance between assembly, inheritance, and environmentally-driven population fluctuations.

### Eco-evolutionary models predict the structure of marine sponge microbiomes

To test whether our evolved host-microbiome *in silico* assemblages recapitulate empirically observed patterns, we compared the diversity and phylogenetic structure emerging from the eco-evolutionary simulations (same as in Fig. 1D) with data from marine sponges. Because sponges are among the earliest lifeforms with complex microbiomes, they provide a natural benchmark for evaluating the early evolution of host-microbe communities [32]. We focused on *low microbial abundance* (LMA) sponges, a group that is reported to have more limited host-control mechanisms compared to the more complex *high microbial abundance* (HMA) species [33], which are evolutionarily newer and derived from ancestral LMA species [34]. The dataset used comprised 562 microbiome samples from 41 LMA sponge species while model samples were taken from the 500 better fitted host individuals at the end of the final generation (the ones with greater *W*_*h*_), divided into 52 species (Fig. 4A; see Methods - Data).

**Fig. 4.**
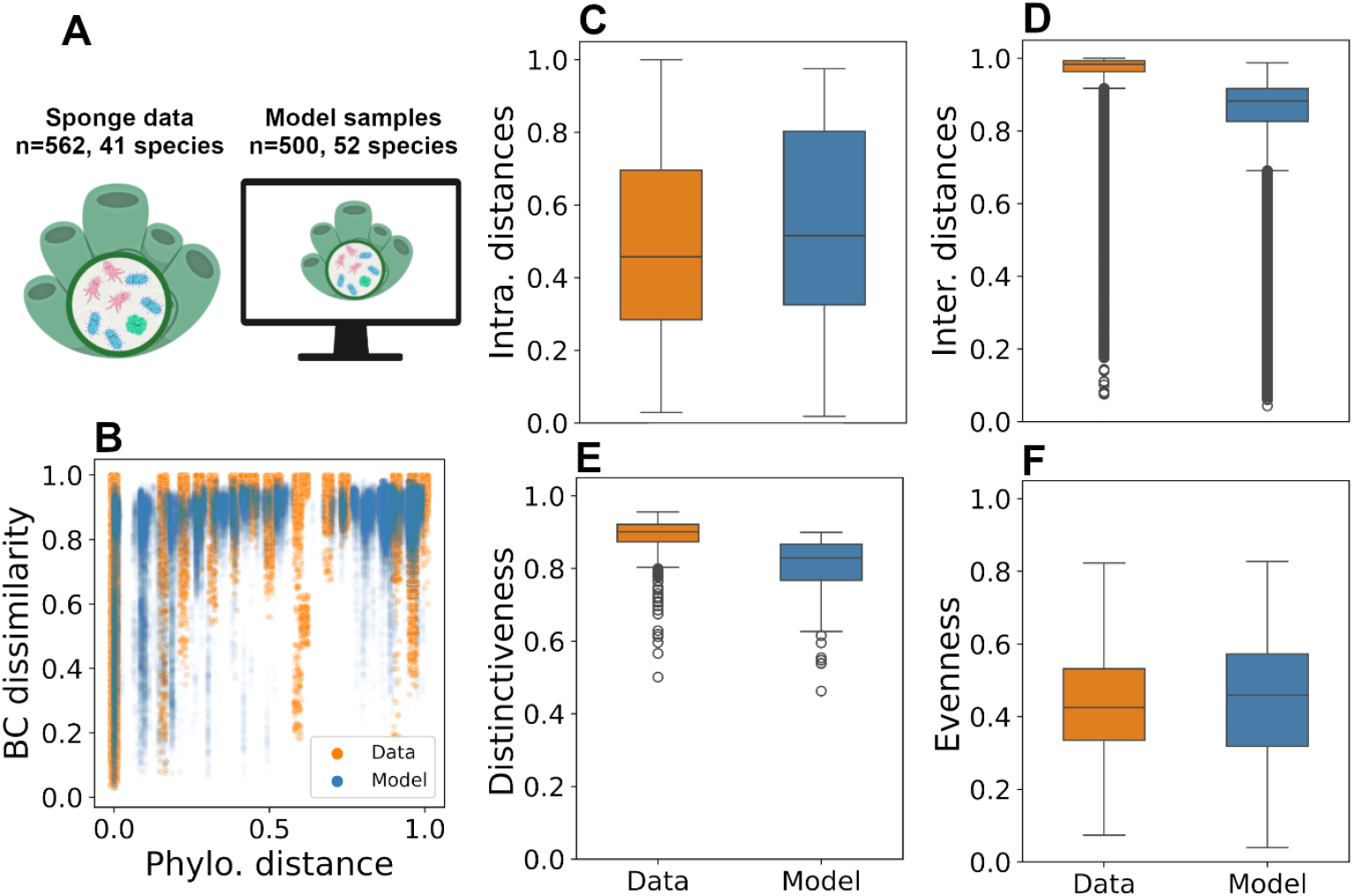
Evolved model microbiomes recapitulate empirical patterns of host-microbe diversity and structure. The panels show comparisons of the final state of simulations (after 2,000 generations; same as Fig. 1D) with empirical microbiome data. (A) Relative microbial abundances from simulations of 500 samples divided into 52 species were compared to 567 samples from 41 species of low microbial abundance (LMA) sponges, organisms expected to harbour largely neutral microbiomes with no or limited host control. (B) Bray-Curtis dissimilarities plotted against normalised host phylogenetic distances. Hosts that are phylogenetically distant exhibit near-maximal dissimilarity, while closely related hosts span the full range of similarity. Empirical data points are less transparent and bigger, shown with stronger jitter, for clearer visualisation. (C,D) Distributions of Bray-Curtis dissimilarity values for (C) within-species and (D) between-species comparisons suggest higher and less heterogeneous dissimilarities between species. (E) Sample-level distinctiveness, measured using a square-root-normalised Simpson’s index for each sample (see Methods -Analysis), shows high values in both model and data, demonstrating that individual microbiomes are typically dominated by a few highly abundant taxa. (F) Distributions of sample evenness (normalised entropy) across model and data are consistent with richly distributed but uneven microbial communities across hosts. All results presented represent the outcome of a simulation starting with *S* = 5 microbial types and evolved to *S* = 103, with generations of 1,000 hosts, sampled in the last generation for 500 best fitted individuals.

We assessed changes in microbiome similarity in relation to host phylogenetic similarity by comparing Bray-Curtis dissimilarities of microbial community structures between pairs of hosts, both in the empirical dataset and in our simulated populations, against normalized phylogenetic distances between hosts. Because Bray-Curtis distances give higher weight to common taxa, this measure highlights the dominant compositional signature of each microbiome. In sponges, host phylogeny was inferred from branch-based distances, whereas in the model it was represented by divergence time, in generation numbers (see Methods – Analysis for details). Both systems exhibited strikingly similar patterns: closely related hosts showed a wide range of microbiome similarities, from nearly identical to entirely distinct, whereas distantly related hosts were very dissimilar (Fig. 4B). This creates a “*forbidden zone*” of moderate to high similarities at large phylogenetic distances, with a sharp transition between variable close relatives and consistently divergent distant hosts.

Within-species comparisons revealed substantial intraspecific variability, with intermediate mean similarity and large standard deviations (Fig. 4C), consistent with the influence of environmental stochasticity. In contrast, between-species comparisons approached maximal dissimilarity with low variance, except for outliers, mostly from closely related hosts, forming a long-tailed distribution (Fig. 4D). Across simulations, results show rich variation with retained consistency of these trends (Fig. S2). For a baseline comparison, neutral evolution, without selection, failed to match the data in all metrics analysed (Fig. S7). These results combined indicate that both in the modelled and in real hosts, microbiomes retain a degree of individuality even under shared ancestry and environment.

To probe the dominance of single (or a few) microbial types in these communities, we computed a square-root normalized variant of Simpson’s dominance index (see Methods - Analysis). Both empirical and simulated microbiomes were highly dominated by a few abundant taxa (Fig. 4E), indicating a skewed composition and a small effective richness. Consistently, evenness (normalized entropy) was intermediate on average but varied widely across hosts, again mirroring empirical observations (Fig. 4F).

Together, these results show that our model recapitulates broad diversity patterns observed within and across host-associated microbiomes. The close agreement between simulated outcomes and empirical observations of community structure in microbiomes suggests that relatively simple eco-evolutionary processes, such as benefits to hosts and vertical inheritance under environmental fluctuations can promote the emergence of natural host-associated microbial communities.

To probe the evolutionary correspondence between host phylogeny and microbiome composition, we compared the structure of host phylogeny with that of microbial community similarity dendrograms, derived from hierarchical clustering of Bray-Curtis dissimilarities. Both empirical and simulated systems showed a lack of phylosymbiosis (i.e. when host phylogenetic distances mirror microbiome distances): the congruence between host and microbial trees was no greater than expected by chance (Fig. 5). The normalized shared phylogenetic information (*nSPI*), which emphasizes early phylogenetic splits, remained low and indistinguishable from randomized controls (*nSPI* = 0.05, *p* = 0.79 for our simulation; *nSPI* = 0.07, *p* = 0.47 for empirical data, 1000 permutations). Similarly, the normalized matching-split information distance (*nMSID*), a complementary metric more sensitive to peripheral tree structure, yielded intermediate values not differing from random (*nMSID* = 0.58, *p* = 0.20 for our simulation; *nMSID* = 0.55, *p* = 0.55 for data).

**Fig. 5.**
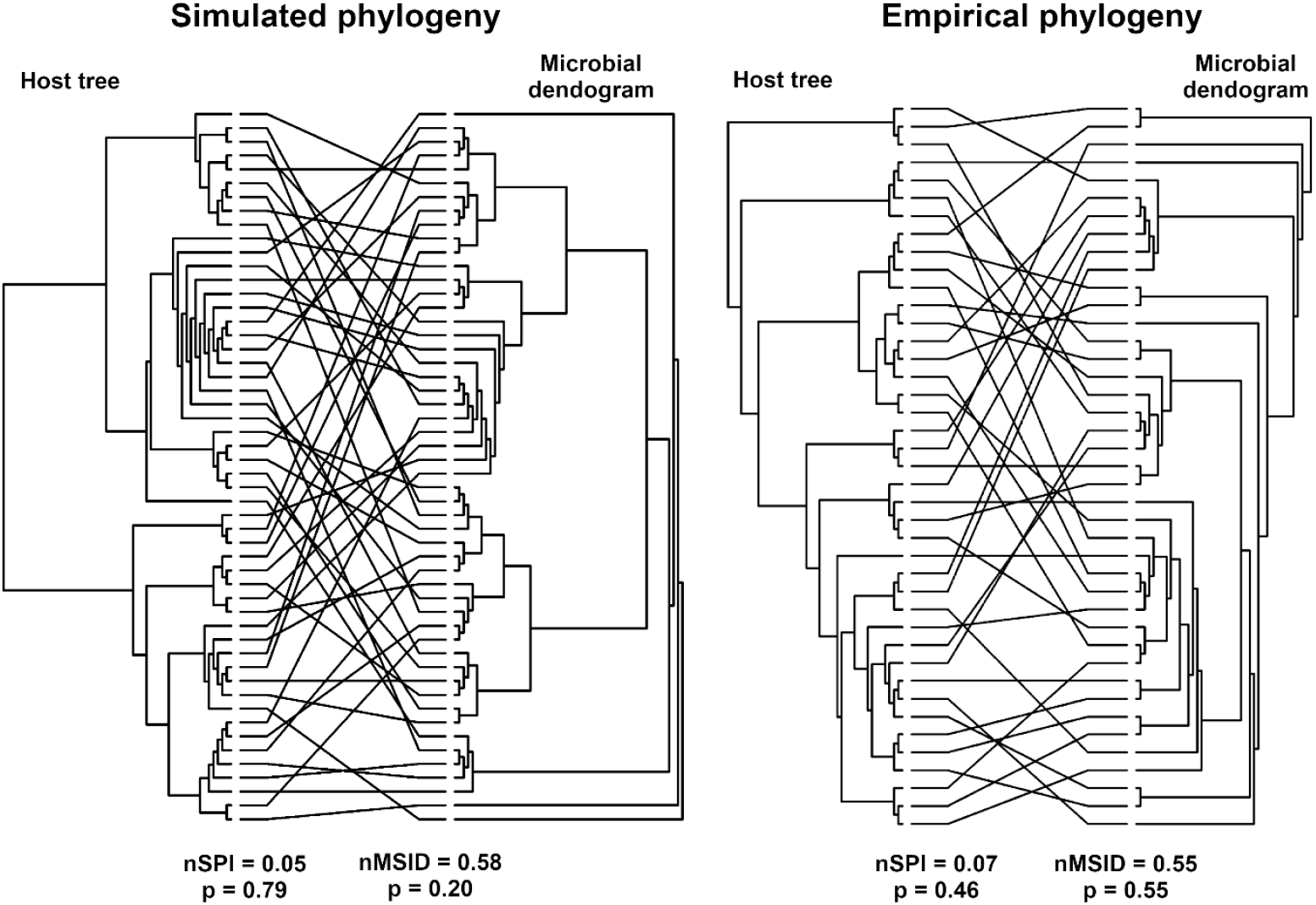
Host-microbiome phylogenies remain incongruent despite the structured evolution of communities. Comparison of host phylogenies and microbiome similarity dendrograms derived from Bray-Curtis dissimilarities. Correspondence between trees was quantified using two information-based metrics that generalise the Robinson-Foulds index while accounting for shared information between trees: normalised shared phylogenetic information (nSPI) and normalised matching-split information distance (nMSID). These metrics emphasise deeper and peripheral phylogenetic splits, respectively. In both cases, observed values were statistically indistinguishable from random permutations (*n* = 1000), indicating no measurable phylosymbiosis. Data is the same as shown in Figs. 1 and 4 for model and empirical communities.

These results agree with the fact that both data and model exhibit a sharp transition in microbiome dissimilarity between closely related and distantly related hosts (Fig. 4B), meaning that microbiome distances directly switch from relatively close to highly dissimilar, without a gradual, more stepwise distancing that follows phylogenetic branching. In other words, an evolutionary relationship between hosts and their microbiomes exists, but it is not strictly encoded as a measurable phylosymbiotic signal. This agreement between model and empirical observations underscores that meaningful structure with an evolutionary fingerprint can arise in host-microbiome systems even in the absence of phylosymbiosis. Mechanistically, this can be a result of the structure of metabolic complementarity, in which species differentiation determines the extent of the matching between a host species and a microbe type. A more intricate complementarity (e.g., one determined by several traits matching) could result in stricter matches and therefore more well-shaped phylosymbiosis. Another important factor is the ability of microbes to survive outside hosts and disperse, thus a more structured microbial flux (*a*) could also result in a higher phylosymbiosis signal.

## Discussion

We developed a mechanistic eco-evolutionary theory of host-associated microbiomes that incorporates three fundamental ingredients: microbial transmission, population fluctuations due to environmental stochasticity, and host benefits. We showed that the presence of beneficial microbes in fluctuating environments, assuming a neutral microbial ecology, can promote the evolution of vertical transmission and the subsequent emergence and persistence of host species-specific microbiomes. Our analyses revealed an inherent homogenising force driven by horizontal microbial immigration, which tends to erode compositional differences across host generations. This tendency can be countered when vertical transmission and stochasticity establish the inheritance and effective selection of microbes. Selection can favour beneficial microbes even when they bear reproductive disadvantages for themselves, and its efficacy is optimal at intermediate levels of environmental fluctuation and host lifespan. The resulting community composition and diversity-phylogeny relationships of evolved microbiomes effectively mirror empirical patterns, reinforcing the robustness of our theory as a hypothesis generating framework for the early evolution of host-associated microbiomes.

Our results outline a mechanistic hypothesis for the initial establishment of beneficial microbes, which contributes towards a description of holobionts in evolutionary theory [35]. In essence, when environmental fluctuations overwhelm minor reproductive differences among microbial types, selection acts primarily through host-level fitness effects. Because microbial taxa can be numerous and functionally diverse, they provide a vast biochemical repertoire from which complementary host-microbe interactions can arise, establishing the beneficial characteristic of some microbes. Environmental fluctuations and indiscriminate transmission are unavoidable factors affecting microbial communities across all environments and hosts. Our model predicts that such plausible conditions are sufficient for the evolution of vertical transmission, which in turn generates enough intergenerational consistency for microbiomes to become inheritable, reliable, and potentially species-specific. However, this is not the only possible route to structured microbiomes. Previous research has demonstrated that complex community structures can evolve even in the absence of host benefits [27], but our work provides a parsimonious starting point for understanding the origins of beneficial associations.

Consequences of these predictions could be tested in future work. For example, experimenters could potentially find a way to control microbial reproductive rates, the magnitude of environmental fluctuations, host lifespan, or the strength of vertical transmission, while tracking the dominance of beneficial microbial lineages. Designed settings with neutral host environments and known beneficial taxa could directly measure the curves of microbial enrichment as functions of microbial reproductive rate, intensity of fluctuations, or host lifespan (Fig. 3). A further hypothesis arising from our results is that hosts might evolve regulatory mechanisms that keep their internal environments within the fluctuation range that maximises microbial selection, effectively harnessing stochasticity to enrich beneficial partners. This might be a simpler, less costly, and more general trait to maintain than any form of specialised host control.

Earlier theoretical work emphasised that maintaining beneficial microbiomes requires a host control strong enough to constrain microbial competition, termed an *ecosystem on a leash* [36]. Our results suggest an alternative scenario by showing that competitive host-specific microbiomes can arise even in the absence of explicit host control, suggesting that beneficial microbiomes can evolve under less restrictive conditions than previously thought. Similarly, Van Vliet & Doebeli (2020) [8] found that strong vertical transmission and short host lifespans are necessary to favour the persistence of beneficial, yet competitively disadvantaged microbes. Our results complement theirs by exploring these requirements across a broader parameter space and demonstrating how vertical transmission itself can evolve in response to environmental variability. The key role of stochasticity that emerges from our model also parallels findings by Bruijning et al. (2022) [25], who highlighted the importance of imperfect transmission for generating host diversity. Although Björk et al. (2019) [37] reported weak and inconsistent vertical transmission in sponges, their analysis focused primarily on the presence or absence of microbe types rather than the relative abundance or functional significance of microbial taxa. This leaves open the possibility that the few dominant or functionally important microbes are transmitted more faithfully, while the majority of rare, peripheral taxa vary randomly, with only a subset of taxa transmitted [38]. In other words, high variability in less abundant taxa may mask consistent transmission of the key microbes that actually shape host benefit. This pattern is fully consistent with the observations of Engelberts et al. (2022) [39] for an LMA sponge species, where more than 90% of the microbiome is composed by three dominant microbe types that are vertically transmitted, providing further support for our interpretation.

The empirical patterns explored in our analysis point to highly species-specific microbiomes yet retaining substantial intraspecific variability. This suggests that host identity exerts a strong influence on microbiome composition, which is consistent with previous observations in host-microbe systems [3,40], while environmental fluctuations likely account for the large variation among individuals of the same species. This pattern is consistent with evolutionary bet-hedging [41], whereby selection favours strategies that reduce variance in long-term fitness under fluctuating environments. With high intra-species variation, some individuals remain close to shifting fitness optima, even though many others are selectively lost. In this sense, variation itself becomes adaptive, contrasting with strict specialisation expected under stable fitness landscapes. Future work might find patterns relating the degree of host specificity to environmental or phylogenetic variables, for example comparing communities in more stable or strongly fluctuating environments. The sharp transition from moderate similarity among closely related hosts to strong dissimilarity among distant ones further suggests that distinct biochemical requirements underpin species-level differentiation of hosts. These observations can be reproduced without invoking additional ecological mechanisms such as active host enrichment of microbes or complex microbial interactions, indicating that general compositional trends and the dominance of few taxa may predominantly arise from the basic eco-evolutionary factors we assumed.

The absence of phylosymbiosis in LMA sponges, as also previously shown by Pankey et al. (2019) [34], provides additional evidence that these microbiomes assemble under strong influence of stochasticity [2] and lack significant host control. This contrasts with observations in HMA sponges, that exhibit signatures of both host control and phylosymbiosis [33,34]. Our theory suggests that hosts may evolve and maintain species-specific, functionally beneficial microbiomes under stochasticity and in the absence of phylosymbiosis and host control. This in turn leads to the hypothesis that microbial function may be a key precursor to the emergence of complex and heritable communities: in microbiomes, function precedes structure. A hypothesis that structure follows function is in line with the widely observed prevalence of functional redundancy in microbial systems [42]. Under this framework, the evolution of host control, microbial genome reduction [12], host-microbe dependence [43], and the eventual signature of phylosymbiosis could represent later stages in the evolutionary trajectory of host-microbe integration, in which deterministic and more structure-dependent processes are more prevalent [2]. Future work can extend our model to incorporate those mechanisms as empirically motivated functions that modulate microbial reproductive rates [9], thus going beyond the assumption of neutrality.

Our results provide a mechanistic foundation for understanding how structured, beneficial microbiomes can emerge from relatively simple coevolutionary processes, prior to the evolution of dedicated host-control traits. Our theory suggests that microbiome evolution may precede, and perhaps even enable, the emergence of more costly enrichment traits and co-diversification mechanisms. In this parsimonious view, the evolutionary history of complex host-associated communities begins with a minimal and plausible establishment of selection and inheritance of microbes.

## Methods

### Simulations

In the simulations shown in Figs. 1, 4, and 5, and in their corresponding replicates (Extended Data), we ran models for 2,000 generations with a population of *H* = 1,000 hosts. At the end of the last generation, the 500 hosts with the highest fitness (*W*) were defined as survivors, and the remaining individuals were discarded.

The initial host generation consisted of a single species with traits *K*_0_ = 5 and *δ*_0_ = 0, and with complementarity *Z*_*h*_uniformly sampled from the interval [0,15]. Each offspring mutated with probability *μ*_*H*_ = 0.01, changing one of its traits (*K*, δ, or *Z*) with equal likelihood to form a new species. Mutation step sizes were drawn as follows:

- For *δ*, from a truncated normal distribution in [0,0.5] with standard deviation 0.1.
- For *K*, from a truncated normal distribution in [1,1000] with standard deviation 5.
- For *Z*, from a normal distribution with standard deviation 1 and circular boundary conditions in [0,15].

Host lifespan was fixed at *T* = 10. The microbial community initially contained *S*_0_ = 5 types. Between generations, a new microbial type appeared with probability *μ*_*m*_ = 0.05, derived from a randomly chosen parent type, with selection probability proportional to the parent’s total abundance across all hosts. Each new microbe was assigned an initial complementarity *z*_*m*_ uniformly sampled from [0,15], and mutation steps from parents were defined analogously to host complementarity mutations. New microbe types were readily added to the external pool, and one host was randomly chosen, weighted by the parent type’s abundance, to receive an addition of 10% of its capacity from the new type’s abundance, before reproduction.

Ecological assembly parameters were held constant as *r* = 1, *a* = 0.1, and *ϵ* = 0.1. In the initial generation, hosts began with empty microbiomes 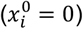. Microbial abundances falling below 10^−6^ were set to zero at every assembly timestep (*dt* = 0.01). We integrated the stochastic differential equations using the Euler-Maruyama method with stochasticity *σ* = 0.1. The host fitness function (*W*) was parameterized with constants *κ*_0_ = 0.5, *κ*_1_ = 10, *γ*_*δ*_ = 0.01, *γ*_*K*_ = 0.001, *W*_0_ = 0.1, and *σ*_*z*_ = 0.5.

For the results shown in Fig. 2C, we used *S* = 100 microbial types, T = 10, *dt* = 0.01, and the same Euler-Maruyama integration scheme. Host traits were fixed at *K* = 100, *δ* = 0.4, *r* = 1, *a* = 0.1, *ϵ* = 0.1, and *σ* = 0.1. A single host species evolved for 100 generations in a population of *H* = 200 hosts, and all hosts were included in the analysis. The fitness function was defined as

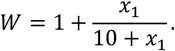

For Fig. 4, the same baseline model was used but with adjusted parameters as stated in the corresponding *x* axes, running *H* = 1,000 hosts for 100 generations. Plots show averages across the last 20 generations, including all hosts.

### Analysis

We computed evenness as the normalised Shannon entropy:

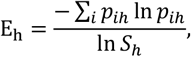

ranging from 0 to 1, where *p*_*ih*_ = *x*_*ih*_/ ∑_*i*_ *x*_*ih*_ is the relative abundance of microbe type i within sample (host) *h* and S_h_ is the total number of types.

The Bray-Curtis dissimilarity between samples *a* and *b* was calculated as:

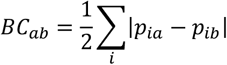

We quantified a diversity index we call distinctiveness (D) as a square-root-normalised measure of the deviation between observed and effective richness:

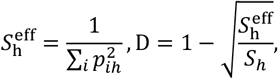

Values of *D* close to 1 indicate microbiomes dominated by a few taxa, whereas smaller values correspond to more evenly distributed communities.

Sponge phylogeny: we derived a species-level phylogeny from taxonomic names using the Open Tree of Life (OTL) database [44] accessed through the rotl R package [45]. Information on divergence times was not retrieved, and phylogenetic distances were qualitatively assessed by counting branches from tip to tip passing through the common ancestor node. Simulated phylogeny: Parent-offspring relationships between species was used to create phylogeny trees and distances were quantitatively measured via species appearance timestamps, in numbers of generations representing branch sizes, instead of just counting branches.

We computed microbial dendrograms by applying UPGMA hierarchical clustering to the Bray-Curtis dissimilarity matrix. To facilitate the visual comparison of host and microbial trees, both were rotated and aligned by matching their tip labels. The procedure iteratively minimizes crossing lines between corresponding taxa by reordering internal nodes while preserving tree topology. We computed two information-based measures of tree congruence (using the *TreeDist* R package). Normalised Shared Phylogenetic Information (*nSPI*) and Normalised Matching-Split Information Distance (*nMSID*) [46-48].

### Sponge data

We analysed host-microbiome associations using data of operational taxonomic unit (OTU), a proxy of a microbial type, from the Sponge Microbiome Project [49]. Sponge species and their low microbial abundance (LMA) classification were obtained from the dataset available in Lurgi et al. (2019) [50], which provided sample identifiers and LMA designations. We focused exclusively on LMA sponges, as their microbiomes are expected to be predominantly shaped by neutral or weakly selective processes, thereby providing an appropriate comparison to our model of limited host control.

Rare OTUs with fewer than 10 total counts across all samples were removed to reduce noise from sequencing artefacts and extremely rare taxa. The resulting dataset comprised 567 samples spanning 41 sponge species. To standardise sequencing depth, we rarefied all samples by random subsampling to 11,331 per sample, corresponding to the smallest number of reads in the dataset. After filtering and rarefaction, the OTU pool contained 122,711 distinct OTUs.

## Data availability

All the data and code used in this study are available in the accompanying Zenodo repository https://doi.org/10.5281/zenodo.17946564.

## Acknowledgements

This project was supported by the Leverhulme Trust through Research Project Grant # RPG-2022-114 to ML.

## Supplementary Information

### Supplementary methods: analytical derivations

We start with the deterministic system for microbiome assembly without microbial turnover (ϵ = 0):

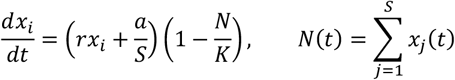

The variable *x*_*i*_ is the density (that we call abundance) of microbe type *i* within the host, with the sum *N* being the total density. Other parameters are constant during the assembly process: *r* is the microbial reproduction rate; *a* is the total influx of microbes from the external environment into the host; *S* is the number of microbe types available; *K* is the host’s capacity (maximum microbiome density). This model assumes that the rates at which microbes are able to reproduce or attach to the host’s internal environment are proportional to the available space within the host. If the community is crowded, it is less probable that new microbes will effectively join the community.

First, we can solve this system for the total density *N*(*t*). We sum the equation for all *x*_*i*_:

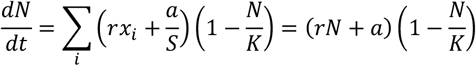

Then we can directly integrate the single equation for *N*. We just need to apply partial fractions to the integral:

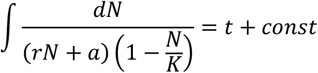

Then we can write the result as a function of the parameter 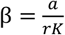:

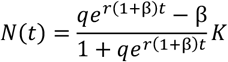

Defining *N*_0_ as the initial density and the constant *q* as:

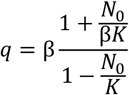

To calculate the assembly time (Fig. S4), we solve for *t* considering that *N*_0_ = 0 and the assembled density defined as a fraction *p* of the capacity *K* (a percentage of assembly). Since *N* asymptotically approaches *K*, it only reaches *N* = *K* exactly with infinite time. However, a *p* = 99% can be quiclky achieved. The final expression for the assembly time is:

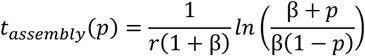

Once we have *N*(*t*), we can use it to directly integrate for each *x*_*i*_:

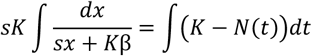

Then, we obtain the exact expression for *x*_*i*_(*t*), with *x*_*i*_(0) being the initial density of the *i* − *th* type:

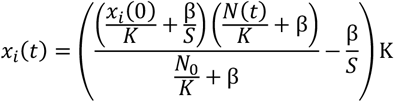

The final equilibrium of *x*_*i*_, with *N* = *K*, is invariant to changes in composition and depends on initial conditions. Therefore, this system preserves a memory of the initial state. This fact highlights the relevance of the vertical transmission mechanism. This equilibrium is:

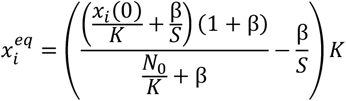

Assuming vertical transmission effectively represented as the offspring’s initial microbiome being a fraction δ of the parent’s final microbiome, we can write the offspring final microbiome as a function of the parent’s final microbiome:

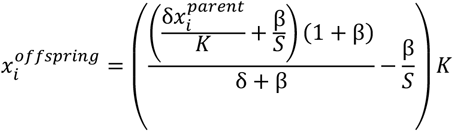

This expression is the same for any type. Then, for any two types *i* and *j* we have:

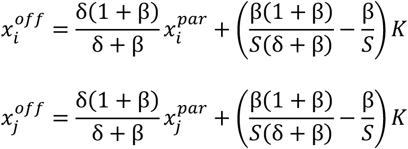

To analyse the convergence of the microbiome towards a state of homogeneity, we can calculate the differences between a pair of species and how they change across a generation. In a way, we can interpret the preservation of structure as the preservation of this distance. By subtracting the two expressions above, we arrive at:

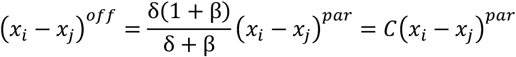

We define *C* as a structure retention factor, measuring precisely the proportion of the difference that is maintained between parent and offspring:

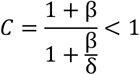

Since *C* < 1 always, it is impossible to perfectly maintain the structure, unless of course that δ = 1, meaning that the entire microbiome is transferred and there’s no assembly dynamics.

By including a microbial death rate ϵ, there is a continuous turnover of microbes. Then, differences in densities are constantly eroded until the only equilibrium of complete homogeneity is reached. However, for small ϵ, the process of assembly can happen with a separation of timescales between a fast assembly and a slow turnover until homogeneity (as seen in Fig. S3). In that case, we can obtain an expression for *C* as a function of the time spent in the slow turnover regime, *T*. As we can also see in Fig. S3, which defines the fast assembly, is that *N* quickly approaches a maximum value. This final *N* can be calculated from the equation for *N*(*t*) by making ⍁{dN]{dt} = 0, solving the resulting quadratic equation for *N*, and ignoring terms of *𝒪*(ϵ^2^):

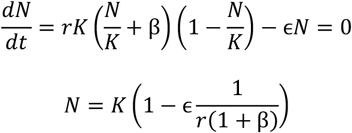

We can see this as a perturbation on the small parameter ϵ moving *N* away from *K*. Then, we write a differential equation for the difference *y*_i*j*_ = x_i_ − *x*_*j*_:

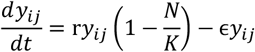

With that, we can obtain the total loss of structure approximately as the loss measured by *C* from before plus an additional loss of structure stemming from spending a time τ in the slow turnover regime, after assembly. The solution of *y*_i*j*_(t) considering the constant perturbed *N* we calculated measures the additional loss:

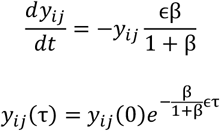

This means that the separation at the end of assembly, which is approximately *y*_i*j*_(0), further diminishes by the exponential factor as the host keeps living. Thus, the combined updated expression for the structure retention factor is:

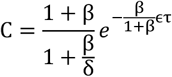

---

**Fig. S1.**
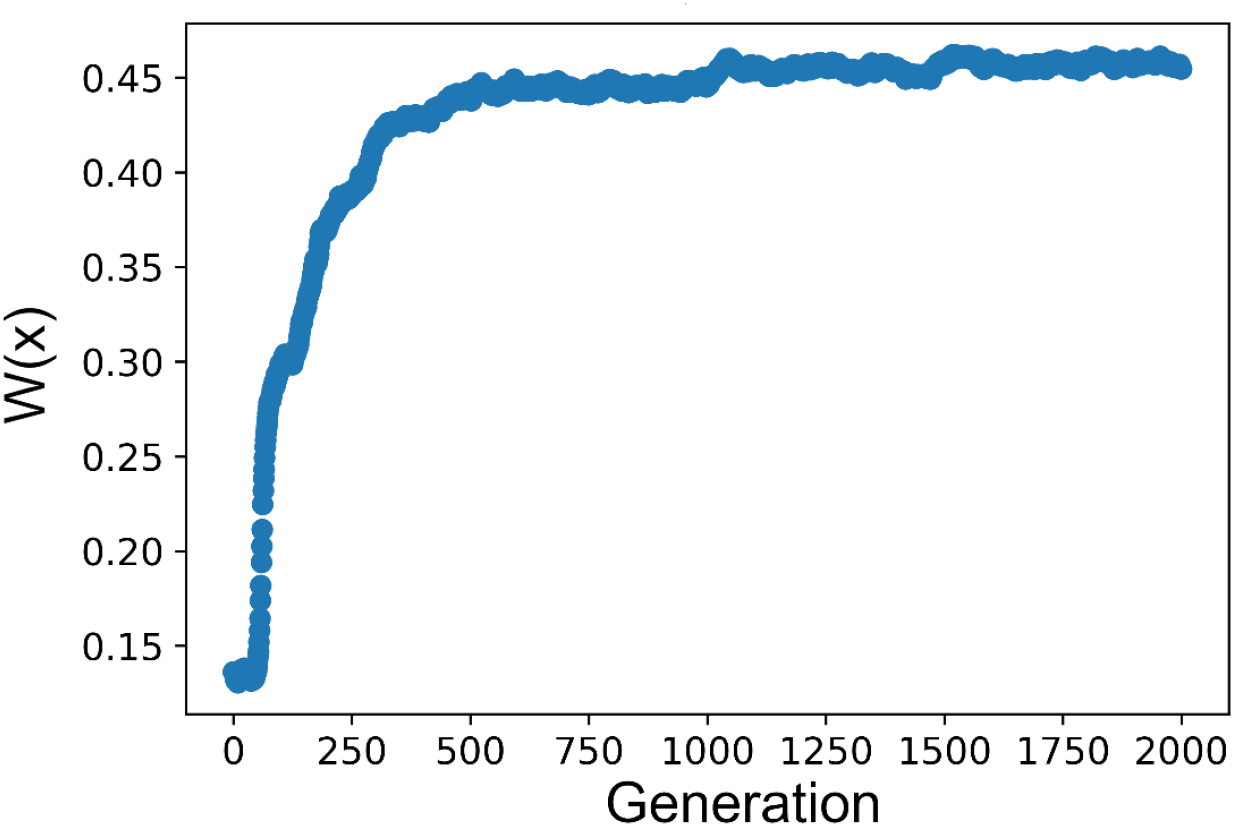
Host fitness for full trait-evolution simulation. Fitness function across generations for the full simulation (*W*(*x*)) presented in Figs. 1D, 4, and 5.

**Fig. S3.**
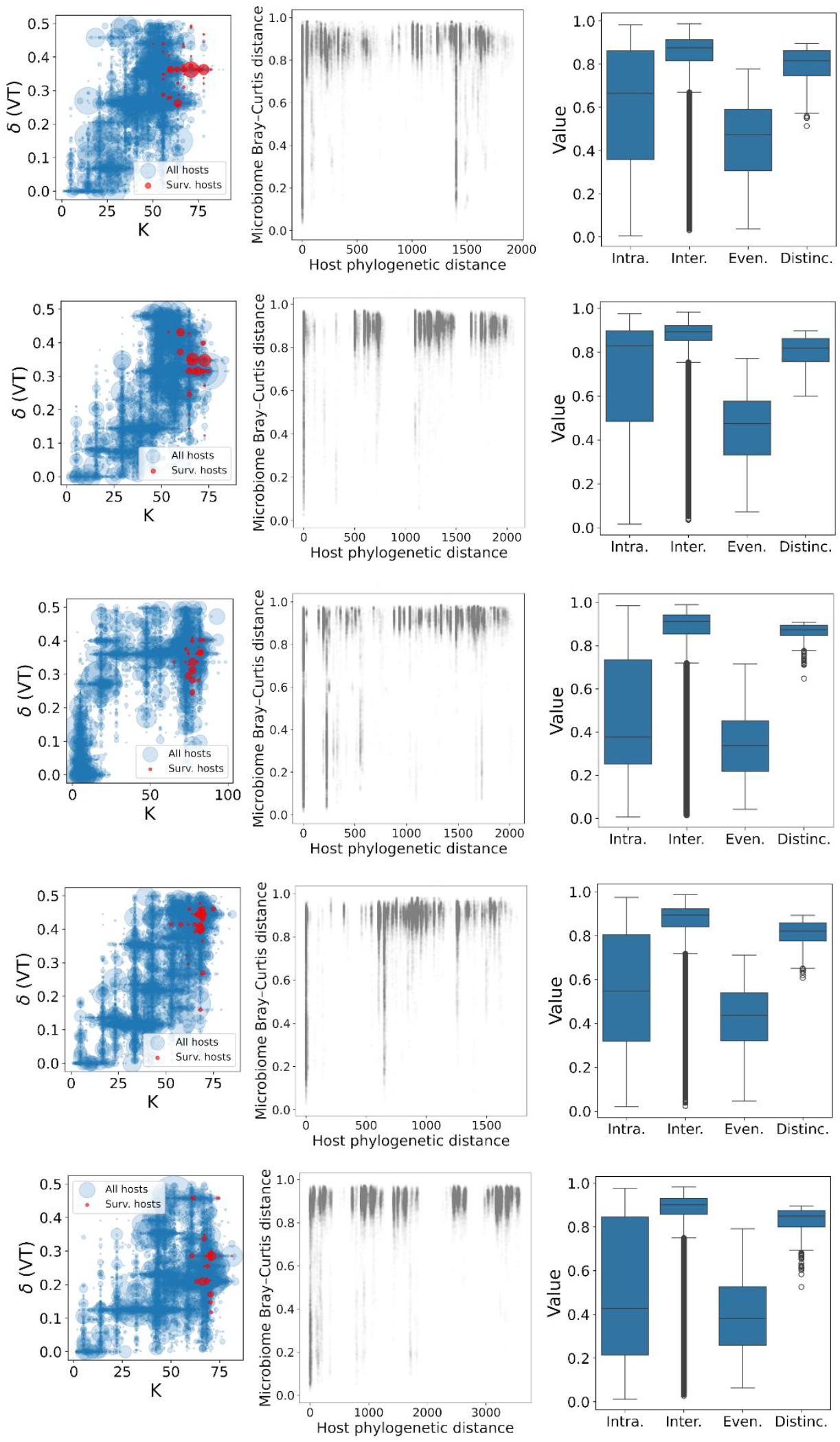
Separation of timescales: fast assembly and slow turnover. Several samples equivalent to the full simulation (*W*(*x*)) presented in Figs. 1D, 4, and 5. Each row shows a sample: (*left*) Evolution of *K* and δ in trait-space, same as shown in Fig. 1D. (*centre*) Dissimilarities by phylogenetic distance, same as shown if Fig. 4B. Empty vertical regions are a result of species extinctions. (*right*) Distances and abundance distribution metrics as shown in Figs. 4C (Intra.), 4D (Inter.), 4F (Even.), and 4E (Distinc.).

**Fig. S3.**
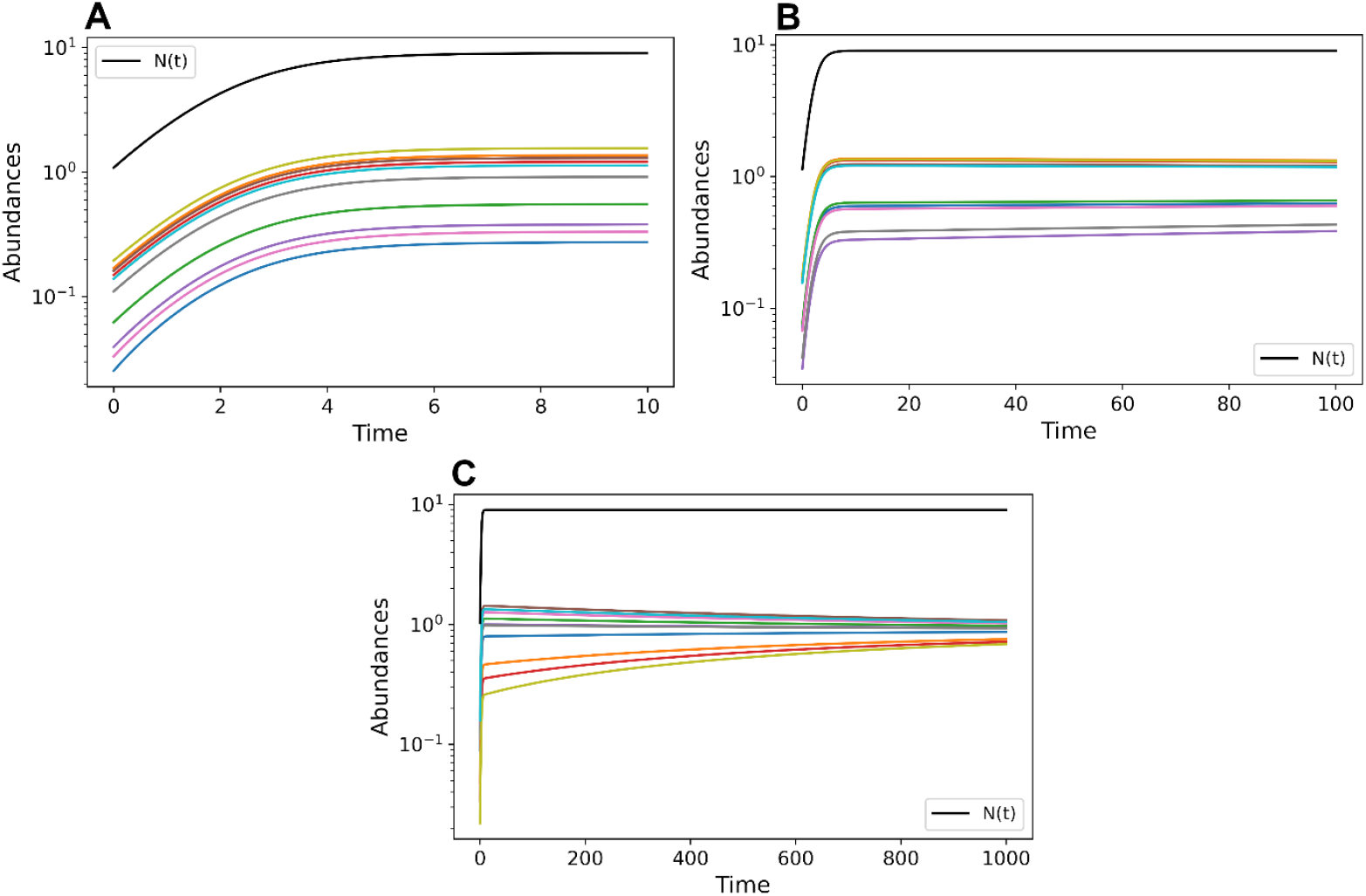
Separation of timescales: fast assembly and slow turnover. Microbial abundances for the deterministic assembly dynamics with *K* = 10, *a* = 0.1, *r* = 1, and ϵ = 0.1, with *S* = 10 microbe types starting from random initial states occupying a total initial abundance of *N*_0_ = 2 (equivalent to the initial microbiome of an offspring with δ = 0.2). Total abundance *N* is shown in black. Plots show the same system but integrated for different durations (10, 100, 1000). (A) The first highlights the assembly regime, with timescale *t* = 10. (B) The second highlights how abundances change in the assembly regime while staying mostly constant in the remaining duration, with the total density changing only during the assembly regime. (C) The third highlights the fact that in the turnover regime, after the assembly and without changes in *N*, individual abundances indeed slowly change towards homogeneity.

**Fig. S4.**
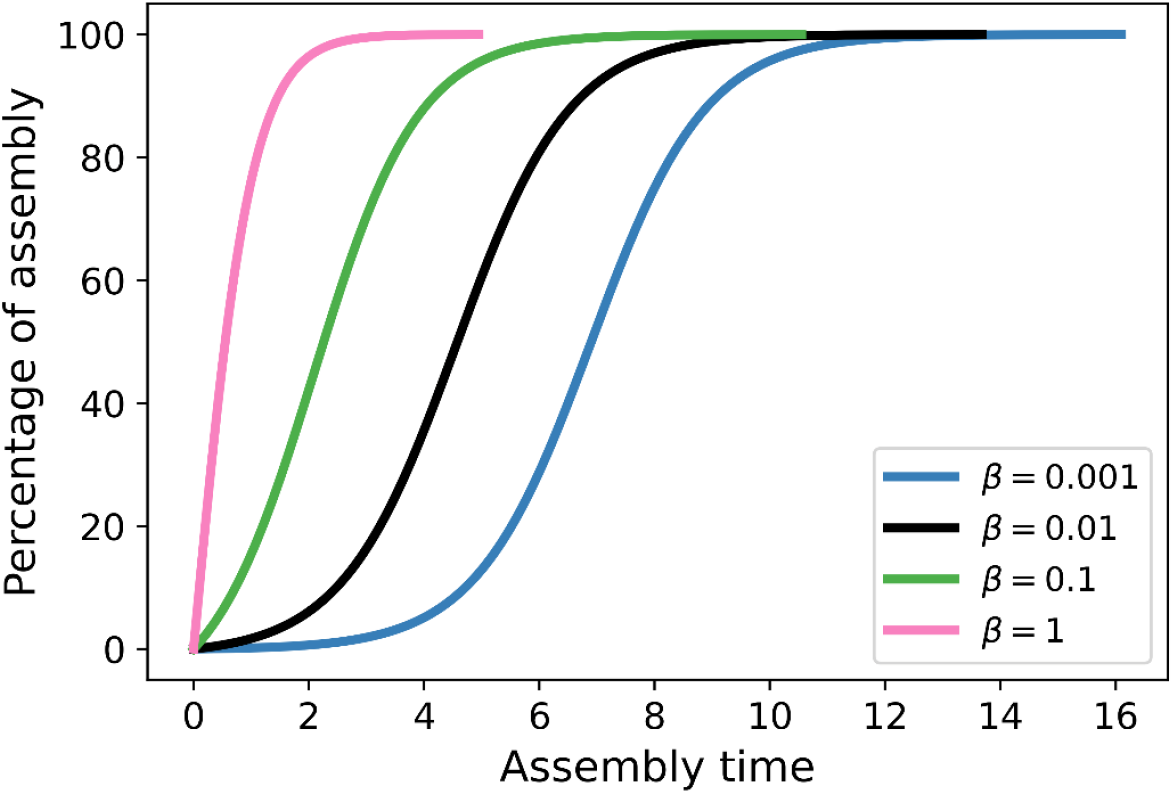
Timescale of microbial assembly. Plot of *p*(*t*_*asse*m*bly*_) according to the function derived in the Supplementary Methods, with *r* = 1. If *K* = 50 and *a* = 0.1, we have β = 0.005, which is approximately what we have in our main trait-evolution simulations. This gives an assembly regime of duration around *t* = 10.

**Fig. S5.**
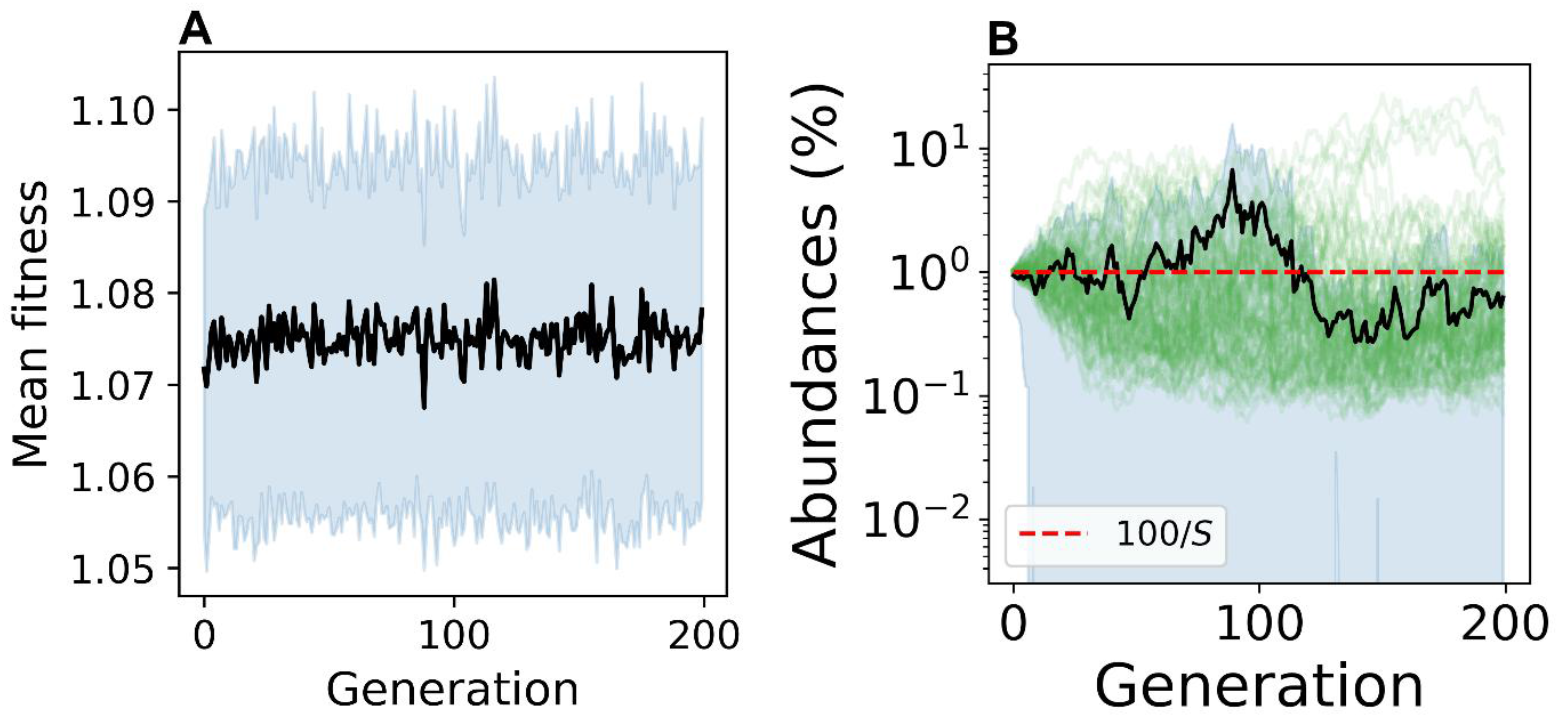
Selection without vertical transmission. Analysis as shown in Fig. 2C for a simulation without vertical transmission, δ = 0, as a baseline of comparison. In this case, we can not expect that beneficial microbiomes can be selected.

**Fig. S6.**
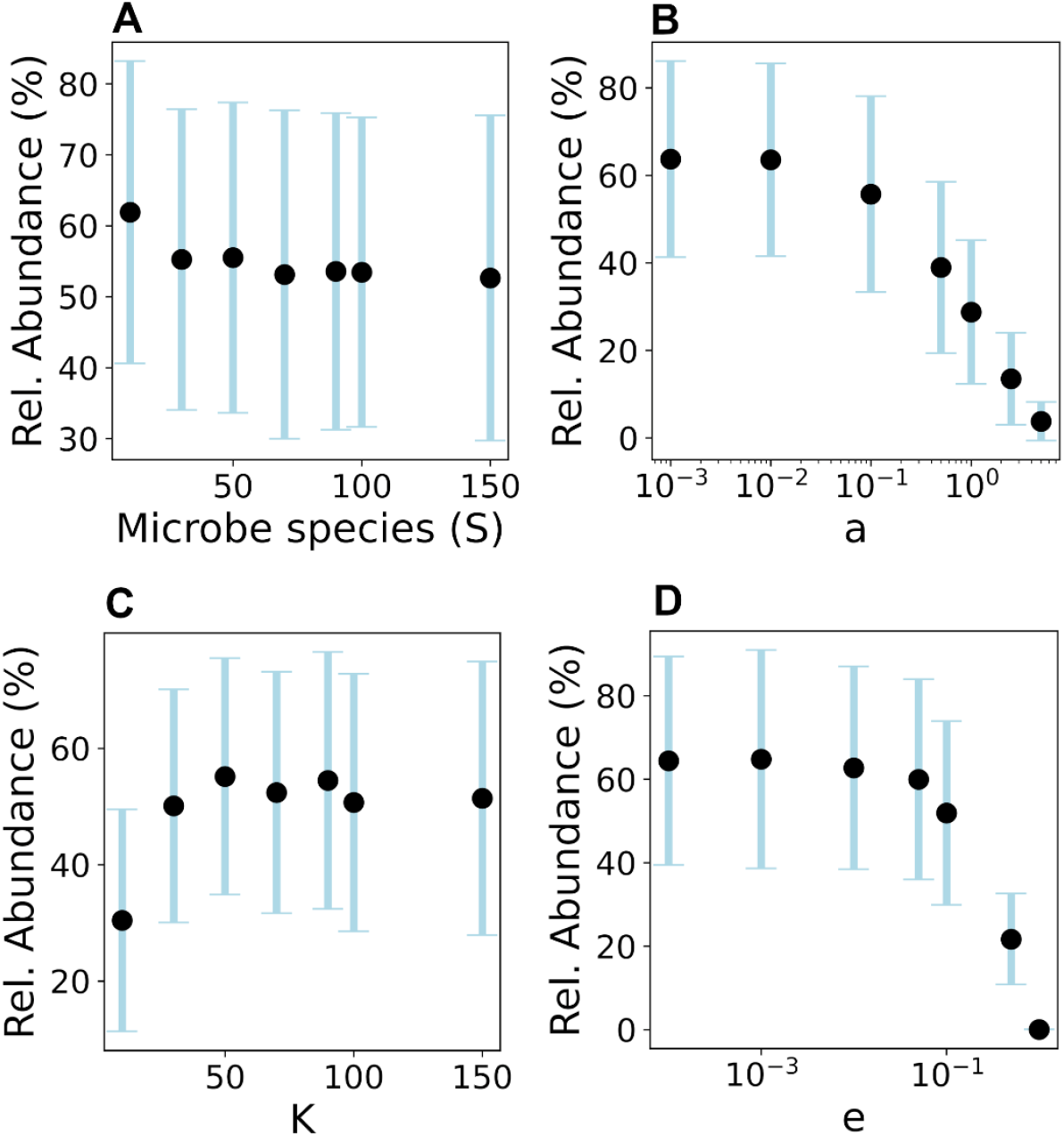
Additional responses of microbiome evolution to ecological parameters. Analysis as shown in Fig. 3, varying: (A) the number of microbe types (*S*), (B) total microbial influx (*a*), (C) host capacity (*K*), and (D) microbial death rate (ϵ). Curves behave as expected: since the total influx of microbes is constant, selection does not directly depend on *S*; it decreases as the total influx increases; it saturates with *K* because of fitness saturation (also visible from the full trait-evolution simulations in Fig. 1D); and it also decreases with the death rate.

**Fig. S7.**
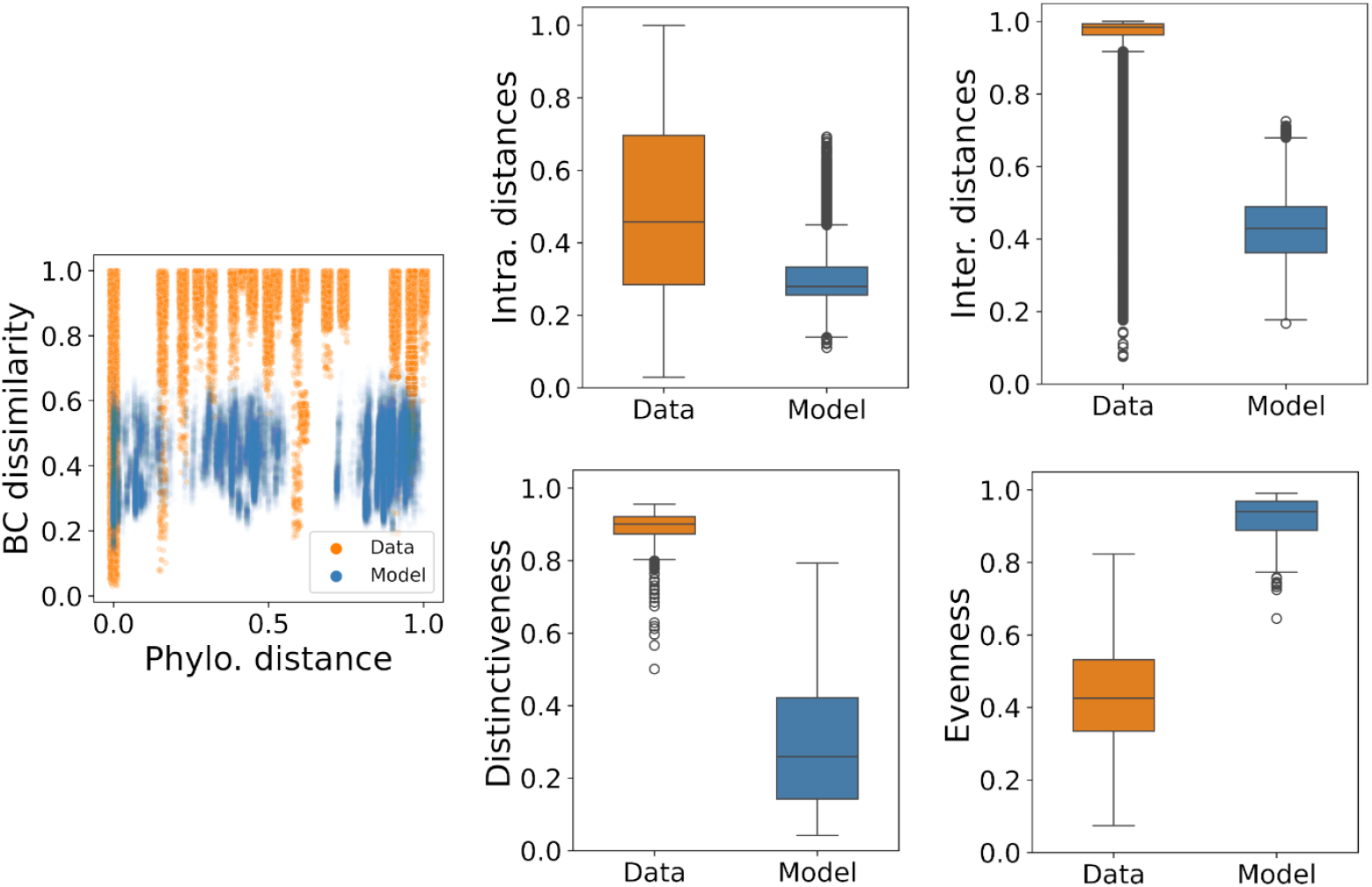
Comparison of dissimilarity and abundance distribution metrics between marine sponge data and the neutral evolution model. Same analysis as shown in Fig. 4, but comparing the data with the simulation without selection (no fitness differences, *W* = *W*_0_). We have 500 surviving hosts after 2000 simulations, here divided into 47 species .

